# Bridging the gap between the connectome and whole-brain activity in *C. elegans*

**DOI:** 10.1101/2024.09.22.614271

**Authors:** Matthew S. Creamer, Andrew M. Leifer, Jonathan W. Pillow

## Abstract

A fundamental goal of neuroscience is to understand how anatomy determines the functional properties of the nervous system. However, previous work has failed to show how the functional connections between neurons are derived from the connectome in the nematode *C. elegans*, raising questions about the extent to which anatomy is informative of signaling^1–3^. Here, we address this problem using a connectome-constrained dynamical model of the brain, which we fit to whole-brain recordings of neural activity during optogenetic perturbation of single neurons^2^. This dynamical model, which contains non-zero weights only between anatomically connected neurons, captured causal interactions between all pairs of neurons 82% as well as the reproducibility of the perturbation data themselves. This included interactions between anatomically unconnected neurons, which the model accounted for in terms of signal propagation over paths that include multiple neurons. Strikingly, alternative models fit using a shuffled connectome achieved much lower performance. Finally, we found that adding connections beyond those in the connectome did not improve the model’s ability to capture causal interactions. Our dynamical model thus provides a link between the connectivity of the *C. elegans* nervous system and its causal interactions and provides a blueprint for exploring the link between structure and function in other organisms.

## Main

A longstanding question in neuroscience is how an animal’s anatomy gives rise to the functional properties of its brain. The nematode *C. elegans* is at the forefront of this question because its connectome is known^4–6^, recordings of whole-brain neural activity at single-cell resolution are available^1–3,7–10^, neurons are stereotyped across animals^11^, and it is an active area of whole-brain modeling^12–21^. However, it has been surprisingly challenging to relate the functional connections measured in neural activity to the anatomical properties of the network in *C. elegans*^1–3^. This leads to a fundamental question: how does the connectome give rise to the functional properties of the brain?

Previous work in this area has shown that including anatomical constraints can allow models to predict neural responses to sensory stimuli^14,22–24^. However, these previous studies did not in general examine causal interactions *between* neurons. Furthermore, past work has been unable to validate predictions of neural-to-neuron interactions against experimental ground truth. This is particularly significant, because several recent studies demonstrate substantial challenges remain in relating anatomical structure with pairwise neural interactions across the brain. For instance, the strength of correlation between two neurons’ activity is poorly correlated with the number of synaptic connections between them^1,3^. Synapse count also correlates poorly with the neural activity evoked by optogenetic stimulation^2^.

These previous efforts highlight the challenges posed by any effort to unify connectivity and signaling. While the connectome provides information about which neurons are synaptically connected, it is unclear how best to infer the weights or signs of those connections in *C. elegans*. Empirical evidence also suggests that neurons signal extrasynaptically, thereby bypassing the connectome, including on short timescales^2,25,26^. Despite this, we still expect the connectome to be critically important for neural signaling in the brain. Here we set aside the known extrasynaptic signaling and instead ask, how well can a model do when considering only the connectome?

Here, we use whole-brain dynamical systems modeling to explain how activity propagating through the connectome of *C. elegans*, including across multiple hops, could generate the observed causal and correlative interactions between every neuron in the brain. In this model, the functional weight between synaptically connected neurons was learned from observing neural activity, while the weight between unconnected neurons was set to 0. This constraint sparsifies the weight matrix such that only 9.4% of possible neuron-to-neuron connections are learned. Because the model is a dynamical system, the neural activity of each neuron is propagated forward to its synaptic partners at every time step, thus bridging the gap between the direct synapses in the connectome and the multi-hop connections that generate the measured neural responses to perturbation.

We demonstrate that this simple linear model captures both causal and correlative structure observed in the data. We further show that adding connections beyond those found in the connectome does not improve model performance, and constraining the model to alternative shuffled connectomes reduces performance. The model predicts that multi-hop signaling is prevalent even between neurons that are directly connected, emphasizing the importance of considering the network effects of the entire brain when modeling neural interactions. Taken together, our model represents the strongest link to date between the brain’s anatomical structure and causal neural responses at single-cell resolution.

### A linear dynamical model of neural activity

We modeled whole-brain neural dynamics in response to optogenetic perturbation using a latent linear dynamical system (**Eq. 1, Fig 1A**). Each neuron’s activity is modeled as receiving a weighted combination of every neurons’ activities on the previous time step, plus its linearly filtered history of optogenetic stimulation:

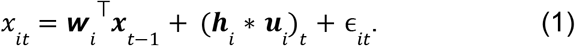

**Figure 1.**
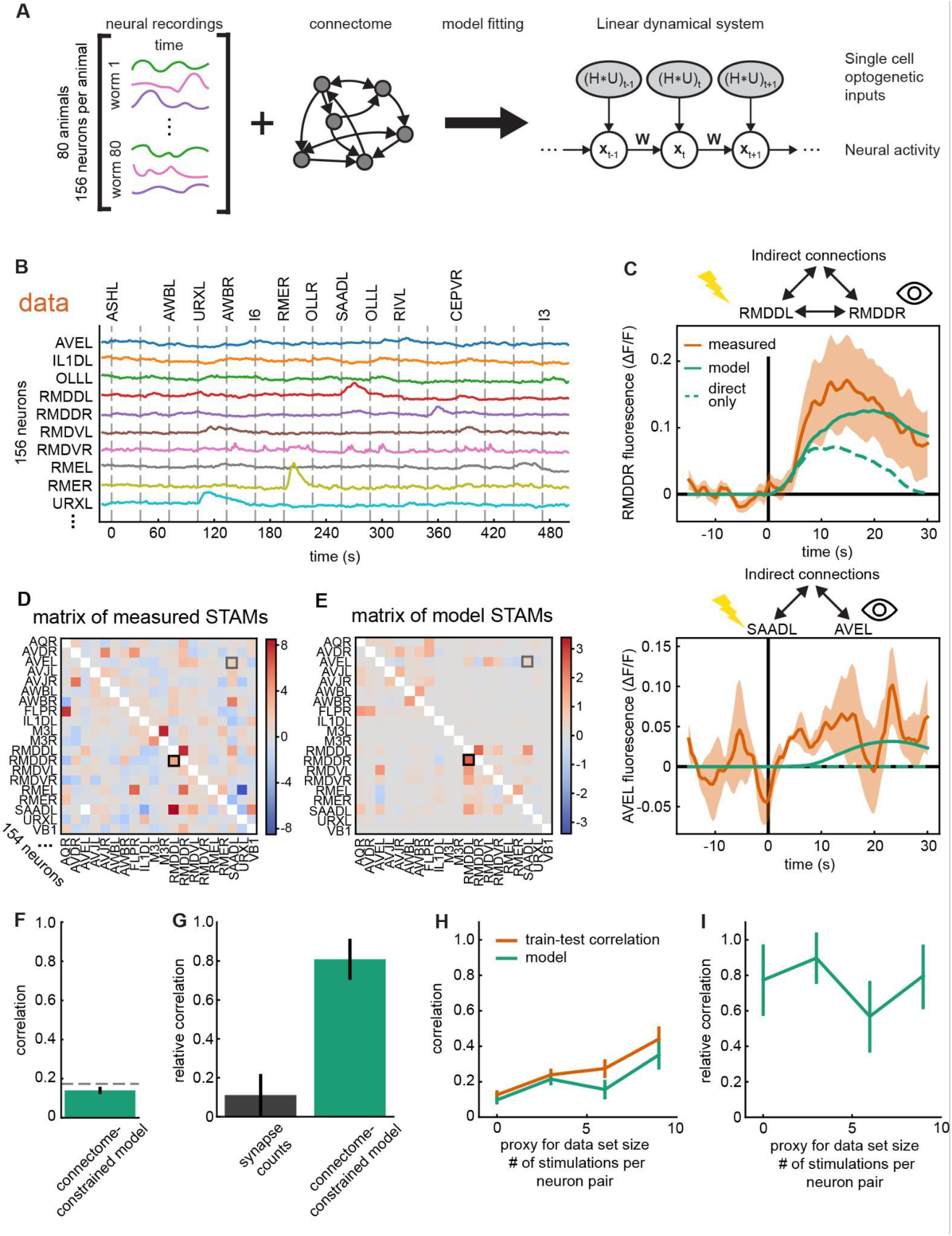
Connectome-constrained model can reproduce the causal structure of whole-brain data. **A**) Diagram of model construction: a linear dynamical systems model constrained to the connectome and fit with whole-brain neural recordings at single-cell resolution. *H* represents the matrix of filters governing the response of each neuron to optogenetic stimulation, *U* is the matrix of each neuron’s stimulation history over time, and *W* is the matrix of neuron-to-neuron synaptic weights. **B**) Calcium activity time traces from 10 example neurons from a whole-brain recording of neural activity. **C**) Top: stimulus-triggered average (STA) of RMDDR activity following optogenetic stimulation of RMDDL (red trace), alongside model-predicted STA using full model (green solid) or a model pruned to contain only the direct connection from RMDDR to RMDDL (green dashed). Bottom: analogous plots showing measured and model-predicted STAs of AVEL activity following stimulation of SAADL, which has no direct synaptic connection to AVEL in the connectome. Error bars show ±SEM. **D**) Matrix of measured STA magnitudes (STAMs) for all pairs of neurons in an example 20-neuron subset; the full data set and model had 154 neurons. Black and gray boxes correspond to the neuron pairs in **C. E**) Matrix of STAMs from the model. **F**) Correlation of the model STAMs to measured STAMs across all neuron pairs (green bar). Dotted line is the train-test correlation. **G**) Relative correlation of the synapse counts and connectome-constrained model to the measured STAMs. Relative correlation is the model’s STAMs correlated to the measured STAMs divided by the train-test correlation (see main text). Synapse counts are calculated as the sum of the electrical and chemical synapses for each pair of neurons and because they lack sign information, they are compared to the absolute value of the STAMs. **H**) Correlation between model STAMs and measured STAMs where the measured STAMs are segmented into pairs of neurons with different numbers of stimulation events as a proxy for data set size. **I**) Relative correlation of the model prediction as a function of data set size. Error bars for **FGHI** are 95% confidence intervals, estimated with bootstrapping.

Here *x*_*it*_ is neuron *i*’s activity at time *t*, ***x***_*t-1*_ is the vector of activity of all neurons on the previous time step, ***w***_*i*_ is a vector of the incoming synaptic weights to neuron *i* from all neurons in the brain, ***u***_*i*_ is the binary time series of laser stimulations of neuron *i*, ***h***_*i*_ is a linear filter representing neuron *i*’s response to optogenetic stimulation, and ϵ_*it*_ is independent Gaussian noise with variance *Q*_*ii*_. Crucially, we set the elements of the weight vector ***w***_*i*_ to zero for all neurons that do not form an electrical or chemical synapse onto neuron *i*. The resulting sparsified model weights contained only 9.4% of all possible pairwise connections. To fit the model, we assumed the neural activity ***x***_*t*_ is not measured directly, but is measured with offset *d*_*i*_ and corrupted by gaussian observation noise with covariance *R* at each timepoint. This leads to a latent linear dynamical systems model, where the neural activity of all neurons can be estimated using the Kalman filter-smoother^27^ (see **Methods**). Modeling the neural activity as latent allowed us to account for missing neurons, as each whole-brain recording inevitably failed to capture a subset of neurons in the brain. This continuous-valued model is well suited to capturing neural activity in *C. elegans*, where the majority of neurons are thought to be non-spiking^28^ and exhibit graded responses^29^.

To fit the model parameters {***w***_*i*_, ***h***_*i*_, *d*_*i*_, *Q*_*ii*_, *R*_*ii*_}, we used recently published whole-brain recordings of calcium activity in *C. elegans* under optogenetic perturbation^2^. These recordings were performed over 20-40 minutes and in total captured 154 neurons in the head of *C. elegans* (**Fig 1B**). The cell identity of each neuron was known^1^, which allowed the model to integrate information from the same neuron across different animals. In this data set, a single randomly selected neuron was optogenetically stimulated with a two-photon laser every 30 seconds. This provided direct estimates of the causal interaction between every pair of neurons in the head, which we used to validate the model. To estimate the average causal interaction for each neuron pair, we computed the stimulus triggered average (STA) of the calcium response in one neuron conditioned on optogenetic stimulation of the other (**Fig 1C** orange). Of the 110 separately recorded animals, we trained the model on an 80-animal training set and evaluated it on a 30-animal held-out test set.

To evaluate the model’s ability to account for this perturbation data, we conducted *in silico* experiments in which we stimulated one neuron at a time and computed the STA of every other neuron, mimicking the *in vivo* optogenetic stimulation experiments. The resulting model STAs accurately captured perturbation responses between anatomically connected neurons, such as the response of neuron “RMDDR” following stimulation of neuron “RMDDL” (**Fig 1C** top green). The model also captured perturbation responses between neurons that were not anatomically connected, such as the response of the neuron SAADL to stimulation of the neuron AVEL (**Fig 1C** bottom green). The model’s ability to capture interactions between neurons that were not anatomically connected relies on the propagation of signals across multi-hop paths through the brain.

To show the magnitude of these multi-hop effects, we additionally calculated the “direct STA” which is an STA generated by a version of the model that only includes the synaptic connection directly between the chosen pair of neurons, eliminating multi-hop signaling. For RMDDR’s response to RMDDL stimulation, we found that both multi-hop signaling and the direct anatomical connection account for a significant fraction of the STA. For neurons SAADL/AVEL the direct contribution is 0 because they do not share an anatomical connection and the entire response is explained by multi-hop signaling.

### Connectome-constrained model accounts for causal interactions in the brain

To quantify the directed causal interaction between each pair of neurons we computed the area under the STA during the 30s after optogenetic stimulation, which we refer to as the stimulus-triggered average magnitude (STAM). We compared the STAMs predicted by the model to the STAMs measured from the neural recordings and found that they had a correlation of 0.14. This correlation is low, but because responses to neural perturbation were variable, the STAMs provide only a noisy estimate of the strength of causal interaction between neurons, which limits how well any model can capture the STAMs. To estimate the variability of the animal itself, we correlated the STAMs measured in the training set with the STAMs measured in the test set (train-test correlation). The train-test correlation was 0.17 and therefore, the model performed nearly as well at predicting the STAMs as the measured data did at predicting itself (**Fig 1F**). To account for the variability of the animal, we report the relative correlation: the correlation of the model to the data normalized by the train-test correlation. While the train-test correlation is not the maximum possible performance of a model, it represents a benchmark where when the model reaches the train-test correlation it is as predictive as the data itself. We found that the model STAMs had a relative correlation of 0.82 to the measured STAMs (**Fig 1G**). We then varied the size of our data set (**Fig 1H**) (see **Methods**) and we found that relative correlation was roughly independent of data set size (**Fig 1I**). The high performance of the connectome-constrained model indicates that the observed perturbation responses can be explained by a network where activity is restricted to propagating along the connectome.

A common hypothesis about the anatomical connectome is that the number of synapses between each pair of neurons correlates with the functional strength between these two neurons. We sought to test this by correlating the absolute value of the measured STAMs with the matrix of electrical and chemical synapse counts from the connectome. The matrix of synapse counts found in the connectome did not correlate with the measured STAMs nearly as well as the connectome-constrained model’s prediction did (**Fig 1G** dark gray). This suggests that the magnitude and sign of the weights learned by the model significantly improve the model’s performance relative to using synapse counts as functional weights.

The STAMs are an estimate of the causal interactions between neurons as measured by perturbation. Another way to measure the interaction between two neurons is by calculating the correlation in their activity over an entire recording. To measure the correlative structure of the data, for each pair of neurons in a recording we calculated the Pearson correlation coefficient of their activity measured over the entire recording. We found that the model’s prediction of the matrix of correlation coefficients provided good agreement to the measured correlation coefficients, thus capturing aspects of both the causal and correlative properties of the brain in a single model (**Fig S1**).

### Model weights differ from synapse counts

We and others have previously observed that the number of synapse counts between two neurons does not directly relate to the strength of the correlation between their activity^1,3^ (**Fig S1B** dark gray) or the strength of their causal interaction^2^ (**Fig 1G** dark gray). This might be expected because synapse counts relate to the direct neuron-to-neuron interactions between neurons while STAMS and correlations are functional measurements that include both direct connections and multi-hop signaling. However, the weights of the model do represent direct connections between neurons, so we wanted to measure how similar the model weights were to anatomical synapse counts. We therefore computed the correlation between synapse counts in the connectome and the absolute value of the model weight for each pair of synaptically connected neurons. Neither chemical (**Fig 2A**) nor electrical (**Fig 2B**) synapse counts correlated strongly with the inferred model weights. Of the two, only electrical synapses showed a modest statistically significant correlation of 0.108. There are two interpretations of this result that are not mutually exclusive. One interpretation is that the model may have found a set of weights that are useful for recapitulating the measurements, but that differs from the weights used by the worm. Another interpretation is that synapse count, as reported from anatomy, may be a poor estimate of the functional strength of neural connections.

**Figure 2.**
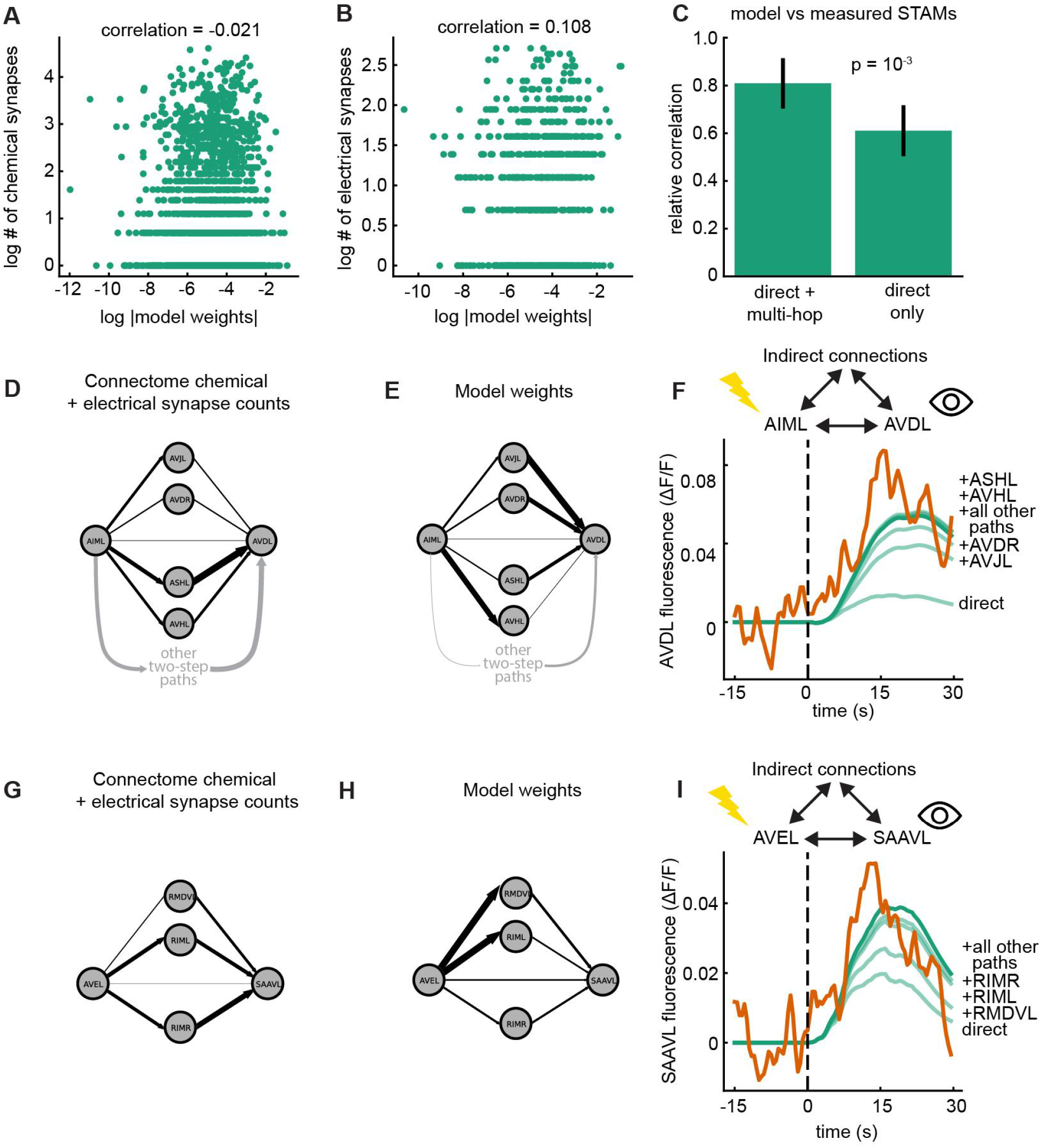
Dynamical model segments neural signaling into direct and multi-hop components. **A**) Scatter plot of log absolute value model weights compared to the log count of chemical synapses between each pair of neurons. **B**) Same as **A**, but for electrical synapses. **C**) Relative correlation between the measured STAMs and the STAMs predicted by the connectome-constrained model (left) and the predictions when the model was constrained to “direct” (monosynaptic) interactions (right). Error bars represent 95% CIs. Error bars and significance calculated with a two-sided two-sample bootstrap test. **D)** Four example feed-forward connections from AIML to AVDL, with arrows weighted by the number of electrical and chemical synapses in the connectome. **E**) Same as in **D** but with the arrows weighted by the magnitude of the model weights. **F**) Comparison between the true STA (orange) and model STAs (green) when different multi-hop paths are enabled sequentially and cumulatively. Dark green represents all synapses activated. **G, H, I**) same as **D, E, F**, except it is the STA of AVAL when AVEL is stimulated.

### Measured causal interactions reflect substantial multi-hop signaling

When one observes an evoked response in the network, that response could include contributions not just from the direct synaptic connection between the stimulated and responding neuron, but all possible paths, including paths that span many synaptic hops through multiple neurons in the network. The model suggests that these multi-hop connections make significant contributions to neuron-to-neuron signaling. We investigated the role of multi-hop signaling by simulating the model’s responses to stimulation, but we prevented signals from propagating beyond the first synaptic partner. When multi-hop signaling was removed in this way, the model performed significantly worse, highlighting the role of multi-hop signaling in the brain (**Fig 2C**). The learned weights of the model presented here provides an estimate of the sign and magnitude of the direct neuron-to-neuron weights after taking into account multi-hop signaling observed in the data. We list the learned weights in a supplemental CSV and table as a resource that researchers can use to compare to other measurements (see **Supplemental Information**).

To find the contribution of multi-hop signaling between two anatomically connected neurons, we considered two pairs of neurons with direct synaptic contacts: AIML/AVDL and AVEL/AVAL (**Fig 2DEGH**). In each case, we found that the evoked response changed substantially when multi-hop signaling was excluded. To investigate this, we silenced *in silico* all the incoming synapses to the postsynaptic neuron AVDL. We then enabled the direct synaptic input from AIML to AVDL and calculated the STA. Since this STA includes no multi-hop signaling, we called this the direct STA. We then sequentially and cumulatively enabled AVDL’s other synaptic inputs in order of their input strength, each time recalculating the STA (**Fig 2F**). In this way we measured the model’s breakdown of how each synaptic partner contributed to the STA of AVDL when AIML was stimulated. We repeated this for AVAL’s response to AVEL stimulation (**Fig 2I**). In both examples, the direct connection was the largest contributor to the STA, but multi-hop signaling contributed a significant fraction of the total response (**Fig 2FI**). We therefore conclude that signal propagation is an important component of the model’s predictive power, and that network effects significantly contribute to causal relationships between synaptically connected neurons. In practice, this suggests that it may be necessary to record from populations of neurons even if one is only interested in understanding how a single neuron influences another.

### Connectome-constrained model outperforms a fully-connected model

Our connectome-constrained model requires signals to propagate along anatomical connections and does not consider extrasynaptic signaling. Does performance increase if we allow the model to incorporate weights between neurons that are not synaptically coupled? To answer this question, we removed the connectome constraint and retrained the model, such that the model learned weights between all pairs of neurons. We then compared the STAMs from the resulting model to the measured STAMs (**Fig 3 AB i**). Despite its significantly larger parameter space, we found that the fully-connected model (**Fig 3AB iii**) performed no better at predicting the measured STAMs than the connectome-constrained model (**Fig 3B v** purple vs green). This result suggests that including connections beyond those found in the connectome does not improve the model’s prediction of the STAMs.

**Figure 3.**
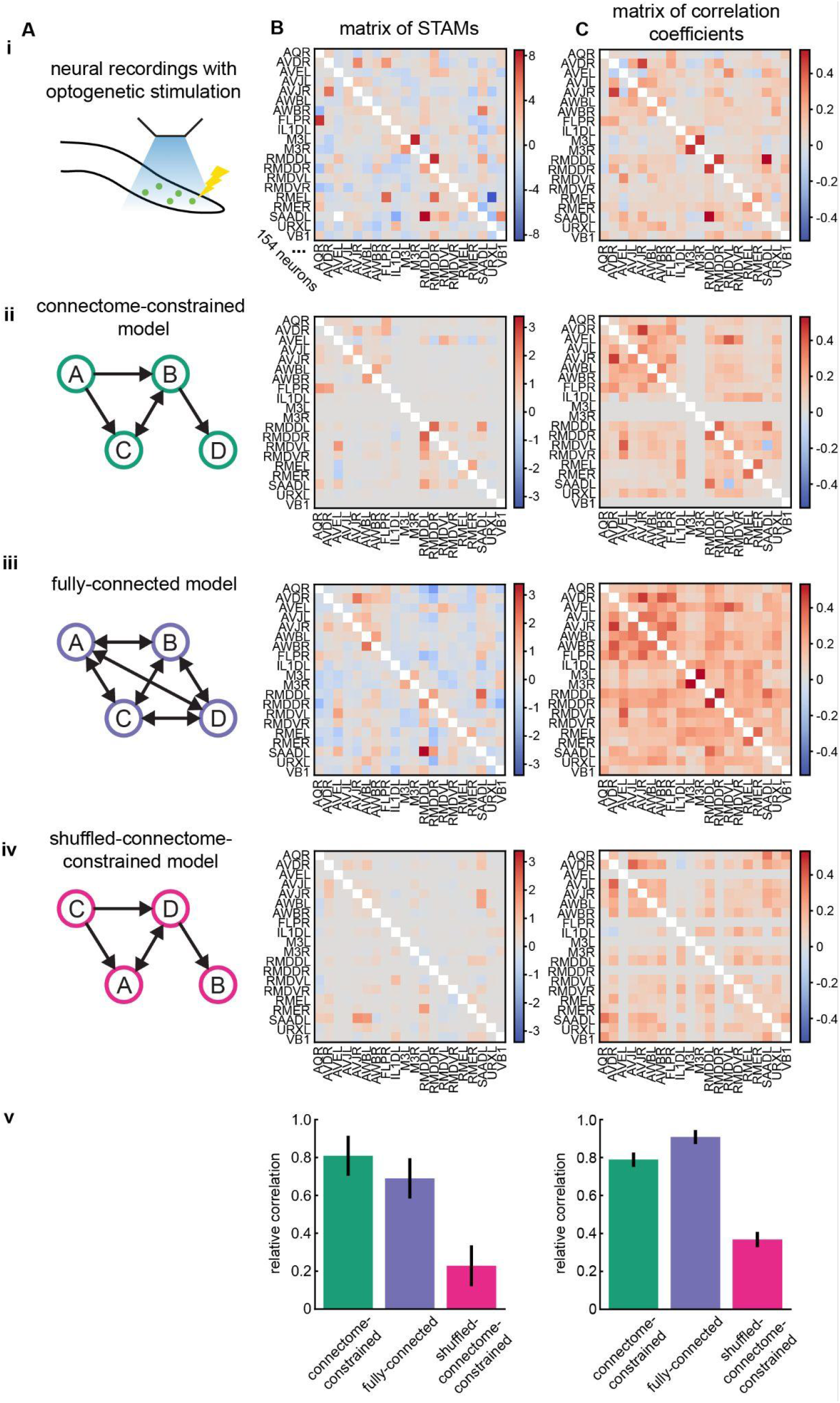
Connectome-constrained model outperforms a fully-connected model in reproducing the causal structure of the data. **A**) Diagram of the data or model in each row. **B**) Matrix of STAMs. **C**) Matrix of correlation coefficients. **i**) STAMs and correlation coefficients measured in the data. **ii-iv**) Prediction of the STAMs and correlation coefficients for each different model. **v**) Relative correlation of the model STAMs to the measured STAMs and correlation coefficients. Error bars are 95% CI, estimated with bootstrapping.

The similar performance between the connectome-constrained model and the fully-connected model is inconsistent with data where extrasynaptic signaling dominates the neural activity. To demonstrate this, we trained a connectome-constrained model on synthetic data generated from a model with significant extrasynaptic signaling. Under these conditions, a connectome-constrained model fails to reproduce the data as well as the fully-connected model (**Fig S2**).

We found that these findings were robust across different model training procedures. Model performance and weights did not change substantially when model parameters were randomly initialized before training (**Fig S3ABC**). We also found model performance was consistent when the model was trained on different subsets of the data set (**Fig S3DE**). Finally, our results are unlikely to be explained by the fully-connected model overfitting: as expected, the test log-likelihood of the fully-connected model was higher than the constrained model (**Fig S3F**).

If adding extrasynaptic connections does not improve model performance, how does one interpret this in light of the known extrasynaptic signaling in the brain? We directly compared performance on 56 STAMs that we had previously identified as relying purely on extrasynaptic signaling^2^. We found that for those 56 neuron pairs the fully-connected model’s performance appears improved compared to the connectome-constrained model though it is not statistically significant on this small set. (p-value 0.1, two-sided paired Wilcoxon signed-rank test, **Fig S4**). Taken together, our findings are consistent with the presence of at least some extrasynaptic signaling in the brain, but including extrasynaptic signaling does not improve model performance when evaluated on all ∼11,000 measured STAMs.

While the extra parameters in the fully-connected model do not improve the model’s ability to predict causal responses on average, they do improve the model’s ability to predict correlations in neural activity (**Fig 3C v**). We calculated the correlation structure of the data by constructing the matrix of Pearson correlation coefficients between every pair of neurons in the head and averaging it over animals (**Fig 3C i**). We then calculated the correlation structure of the brain as predicted by each model (**Fig 3C ii-iv**). We found that the fully-connected model slightly outperformed the connectome-constrained model at capturing the correlation structure of the data (**Fig 3C v**). This result makes sense given that the model is trained to recapitulate the activity traces, but it is not explicitly trained to prioritize predicting STAMs over other aspects of the activity.

### The connectome itself is critical for model performance

We wondered whether the success of the connectome-constrained model at predicting perturbation responses should be attributed to the connectome itself or rather to the sparsity that the connectome provides. For instance, there may be many sparse constraints that allow the model to capture the STAMs but have little to do with the connectome. To test this, we constructed a third model where we constrained the dynamics matrix *W* to the connectome, but shuffled the neural identities (**Fig 3 A iv** note the label changes, see **Methods**). We fit this shuffled-connectome-constrained model to the animal’s neural activity and calculated its predicted STAMs (**Fig 3B iv**) and matrix of correlation coefficients (**Fig 3C iv**). The shuffled model performed poorly at capturing either causal (**Fig 3B v** and **Fig S3** pink) or correlative interactions between neurons (**Fig 3C v** and **Fig S3** pink), suggesting that the model’s performance is due to the specific connections found in the animal. This result was consistent across multiple different shuffles of the connectome (**Fig S3**). Taken together, these results demonstrate the significant role that the connectome plays in producing the observed neural activity.

### Connectome-constrained model predicts the activity of held-out neurons

A fundamental challenge when recording from neural populations is that for any individual recording one typically does not observe all neurons. However, information about the activity of missing neurons is encoded in the network via interactions with their synaptic partners. We found that our model was able infer the activity of unmeasured neurons in the recording using its estimate of the functional connections between neurons.

To assess the model’s accuracy at predicting missing neurons, we held out a neuron’s activity from a novel animal. We then compared the model’s prediction of the neuron’s entire time series against measured ground truth (**Fig 4A**, see **Methods**). Across all held-out neurons and recordings, the model’s prediction of the held-out neuron’s time series achieved a mean correlation of 0.25 to the true activity, significantly higher than a shuffled control in which the prediction was compared to incorrect neurons (**Fig 4B**). The model predicted the held-out neuron AVER the best of all neurons with a correlation of 0.93 (**Fig 4C**). The neuron with the median prediction score was URYVL with a correlation of 0.23 (**Fig 4D**). The model’s ability to reconstruct held-out neurons demonstrates that the model has learned the functional properties of individual neurons that are unique to each neuron and conserved across animals. Notably, our model found this conserved structure in the language of individual neuron-to-neuron connections without relying on a low-dimensional latent space that inherently mixes contributions from many neurons.

**Figure 4.**
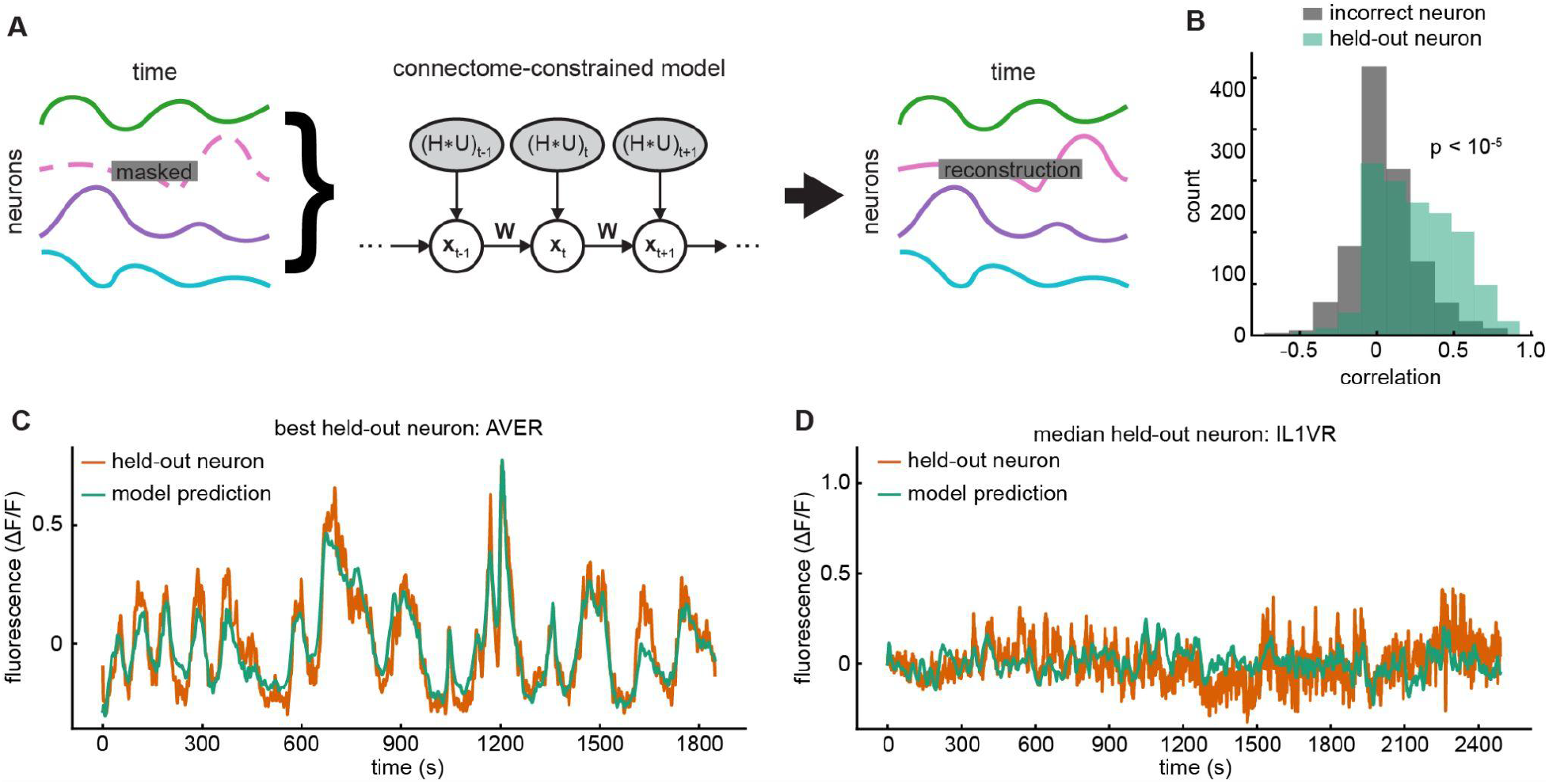
Connectome-constrained model predicts the activity of held-out neurons. **A**) Diagram detailing the process of neuron reconstruction. An individual neuron in a whole-brain recording from a held-out animal is masked such that the model does not have access to it. Then, based on the model’s prediction of the dynamics of the held-out neuron and the activity of the other neurons, the model reconstructs the missing neuron. **B**) Accuracy distribution of the model’s reconstruction where each accuracy score comes from reconstructing a single neuron in a single held-out animal (green). Accuracy distribution of the model’s reconstruction when compared to an incorrect neuron (gray). P-value calculated with a two-sided two-sample bootstrap test. **C**) Example trace of the model’s reconstruction of AVER. **D**) Example trace of the model’s reconstruction of IL1VR.

## Discussion

Synaptic communication is considered to be the dominant method of signal transduction in nervous systems. Despite this, previous work has found a surprisingly weak relationship between measured functional interactions and the anatomical connectome of *C. elegans*^1–3^. Here, we present a dynamical systems model that provides an explanation of how the causal and correlative properties of the brain can arise from signals propagating along a connectome. These findings highlight the importance of dynamical systems analysis for bridging the gap between the connectome and neural function.

The work presented here shows the significant role that the anatomical connectome plays in generating neural activity. We found that our connectome-constrained model captured many of the observed perturbation responses, and adding an order of magnitude more connections outside of the connectome did not improve performance (**Fig 3B v**). Furthermore, constraining the model to a shuffled version of the connectome abolishes the model’s ability to capture both the causal and correlative structure of the data, clearly demonstrating the importance of the connectome in generating whole-brian activity (**Fig 3 v**).

### The role of extra-synaptic signaling

How does our work coexist with previous evidence of extrasynaptic signaling in the brain? Our results do not preclude the existence of known extrasynaptic signaling^2,31^. First, the STAMs reported here are only one measurement of the properties of the nervous system and exclude important functions such as neuromodulation. It is also possible that the success of the connectome-constrained model in explaining these STAMs may come from the flexibility to recapitulate some extrasynaptic signaling by recasting it as synaptic or multi-hop signaling.

Additionally, the significant animal-to-animal variability found in the data (**Fig 1FH** and **Fig S5**) could be masking the role of extrasynaptic signaling. Models trained on an even larger data set could reveal roles for connections outside those found in the connectome. Finally, the fully-connected model performs slightly better for a group of 56 previously identified instances of purely extrasynaptic signaling (**Fig S4**), which is consistent with extrasynaptic signaling in these neuron pairs. Therefore we suspect that extrasynaptic signaling is still present, even though it does not improve model performance when evaluated across all ∼11,000 neuron pairs.

### Relationship to previous work

Our work builds on a rich history of research seeking to use dynamical systems to account for the nervous system’s functional capabilities. Research using dynamical systems to describe whole-brain activity often incorporates either connectomic data^17,18,22,32–34^ or neural data^13,35–38^ but not both. Our approach bears the greatest similarity to work that sought to incorporate both anatomy and neural recordings into functional models of neural activity^14,39^. Our work is distinguished by combining both neural recordings and anatomical data to provide a model of how neuron-to-neuron causal and correlative interactions result from anatomical structure.

### Limitations and future directions

Despite the successes of the model presented here, there are still important caveats to consider. The data used in this work only measured pairwise causal interactions, which does not account for nonlinear interactions within groups of neurons. One potential way to measure these interactions would be to record whole-brain activity while activating multiple neurons using patterned light. From a modeling perspective, the linear dynamical systems like the one we employed are restricted to a single fixed point, precluding phenomena like bistability and limit cycles. These restrictions limit the model to simpler dynamics and underestimates the magnitude of the responses compared to those measured in the recordings. Furthermore, because the model is linear and time-invariant, there are many fundamental properties of the nervous system that the model cannot account for, such as learning, neuromodulation, and state-based processing. These are of particular interest because they may help explain the large variability observed in the perturbation responses (**Fig S5**). Finally, our recordings only measure calcium activity in the neurons, which is only a proxy of the true neural activity. It is possible that the underlying voltage dynamics of the neurons is propagating through the network much faster than we observe in the calcium. We focus on capturing the calcium data we have access to, but future work could attempt to infer the latent voltage dynamics or attempt to measure the voltage dynamics experimentally with genetically encoded voltage indicators.

An intriguing future direction of this work would be to extend the model to incorporate more realistic biophysical properties, such as response rectification, saturation, and nonlinear conductances. The resulting model could exhibit nonlinear dynamics and account for a wider range of biophysical phenomena. Formally, our model can be considered a linear approximation to the dynamics of the *C. elegans* brain, and provides a tractable starting point for the development of more complicated nonlinear models.

## Acknowledgements

The authors acknowledge funding from the Simons Foundation (SCGB AWD543027) and the BRAIN Initiative (9R01DA056404-04).

## Methods

Models were defined and fit using custom python code, available here https://github.com/Nondairy-Creamer/Creamer_LDS_2026 All data used is published^2^.

### Model

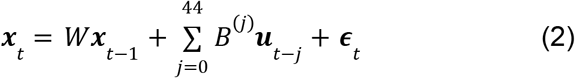

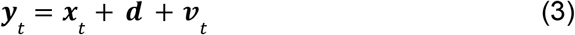

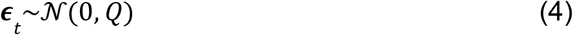

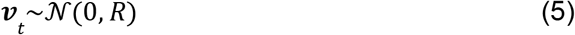

In this model, ***x*** is a latent vector of the calcium activity of every neuron in the nematode brain at time *t. W* is the dynamics matrix, which specifies the direct connection between neurons and is the weights ***w***_*i*_ stacked as row vectors. The neural activity at each time point is approximated as a linear transformation of the neural activity at the previous time point by the matrix *W*. Crucially, the weights in *W* are constrained such that weights between neurons that do not share a chemical or electrical synapse are set to 0, while weights between synaptically connected neurons are learned. ***u***_*t*_ is a one-hot input vector that designates which cell is optogenetically stimulated at time *t. B*^*(j)*^ is a diagonal weight matrix that describes how a neuron responds to being stimulated at *j* timepoints in the past. By enforcing *B* to be diagonal, we ensure that the laser stimulation is only allowed to affect a single neuron and the effect of stimulation on the network must be explained by signal propagation from the targeted neuron through *W*. The matrices *B* are related to the filter ***h***_*i*_ from **equation 1** in that 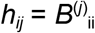 and *h* is 45 timepoints long (22.5 seconds at 2 Hz). The variables ***ϵ*** and ***v*** are vectors of additive Gaussian noise in each neuron with diagonal covariances *Q* and *R* respectively.

***y*** is the observed calcium activity and ***d*** is the vector of offsets between the latents and observations. Not all neurons are measured in each recording, therefore ***y*** is partially observed. By separating our model into the partially observed activity ***y*** and the latent activity ***x*** we can infer the latent activity of all neurons in the brain even when they are not measured in an individual recording. Notably, we perform no dimensionality reduction in the model. This means that every connection in the model is neuron-to-neuron, which allows us to individually perturb each neuron in the model. The latents ***x***_*t*_ were estimated using Kalman filtering-smoothing and the parameters were fit using expectation maximization^27,40,41^. The learned parameters in the model are *W, B*^*(j)*^, *Q, R*, and ***d***.

### Relative correlation

Relative correlation is a ratio between model performance and an estimate of data’s correlation to itself. Model performance is calculated as the correlation between the model’s predicted STAMs for every neuron pair in the head of the animal. Our estimate of data’s correlation to itself is the train-test correlation, which is calculated by correlating the STAMs measured in the training set with the STAMs measured in the test set. The intuition is that if the model perfectly captures the data in the training set, then model performance will be equal to the train-test correlation and achieve a relative correlation of 1. We note however, that this does not imply that the train-test correlation is not the maximum possible performance could achieve. If there is low mismatch between the brain and the model, the model can denoise predictions and achieve higher performance than the train-test correlation. Furthermore, a model that takes into account the state of the animal at the time of stimulation could also outperform the train-test correlation.

### Connectome

For both the connectome-constraint and synapse counts the connectome we used was the sum of the matrix of synapse counts from three adult connectomes^4,6^ and one L4 connectome^4^. All connectomes were taken from https://www.nemanode.org/ on November 1st, 2025. The specific CSVs used were white_1986_jsh.csv, white_1986_n2u.csv, witvliet_2020_7.csv, and witvliet_2020_8.csv.

### Proxy for data set size

In **Fig 1HI** we construct a proxy for data set size. In the perturbation data, neurons are randomly selected for optogenetic activation. This leads to wide variability in the number of times each neuron is optogenetically stimulated and thus how reliably its functional connection to other neurons can be measured. We use this variability by segmenting all possible neuron pairs into different data sets based on the number of stimulation events observed for each pair. In this way, one data set will consist exclusively of pairs of neurons with 1-3, 4-6, 7-9, 10-12, or 13-15 stimulation events. We note that the model’s correlation to the data increases as the number of stimulation events per neuron pair increases, but the relative correlation remains approximately constant (**Fig 1HI**).

### Direct STAMs matrix

This matrix is the model’s reconstruction of the measured STAMs if signals were not allowed to propagate over multiple synapses. To construct this matrix, we individually consider each pair of neurons in the head where neuron A is the stimulated neuron and neuron B is the recorded neuron. We then take the model’s weight matrix *W* and set every weight to 0 except for the weight from A to B, as well as the two neuron’s self terms. We stimulate A and measure the STAM in B. Because all connections to neurons other than B are set to 0, no multi-hop signaling is possible. As a result, any pair of neurons that are not anatomically connected will have a direct STAM of 0 (**Fig 1C** bottom).

### Shuffled-connectome constraint

We constructed this shuffled connectome by taking the binary connection matrix of the connectome and permuting its rows and columns. In this way any given neuron is assigned the connectivity pattern of another randomly chosen neuron in the data set. The resulting network is topologically identical to the original network; we have only permuted the identity of each node in the graph. As a result, the model has to explain the same neural activity with an entirely different set of connections. Once we have this shuffled connectome, the model is constrained in the same manner as the connectome-constrained model: all weights between synaptically connected neurons are learned while weights between neurons that do not share a synapse are set to 0.

### Neuron reconstruction

To calculate the model’s performance on neuron reconstruction we looped through every animal and every neuron in the held-out test set. For each neuron ***x***_*i*_ that was recorded, we masked that neuron’s activity such that the model treated it as missing. We then calculated the posterior of the neural activity of the masked neuron ***x***_*i*_ given the activity of all the recorded neurons at every time point 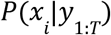 and use the mean of the posterior to estimate the missing neural activity.

Most neurons in the worm brain are bilaterally symmetric and can be highly correlated with their bilaterally symmetric partner. To exclude the possibility that the model trivially uses this information for reconstruction, we additionally held-out the neuron’s partner for each bilaterally symmetric neuron.

### ΔF/F calculation

The ΔF/F reported is calculated with the standard equation 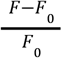 where F is the GCaMP fluorescence data. However, we define F_0_ to be the mean over time of F. As such the units can also be interpreted to be fold-change relative to the mean.

## Supplemental Figures

**Supplementary Figure 1.**
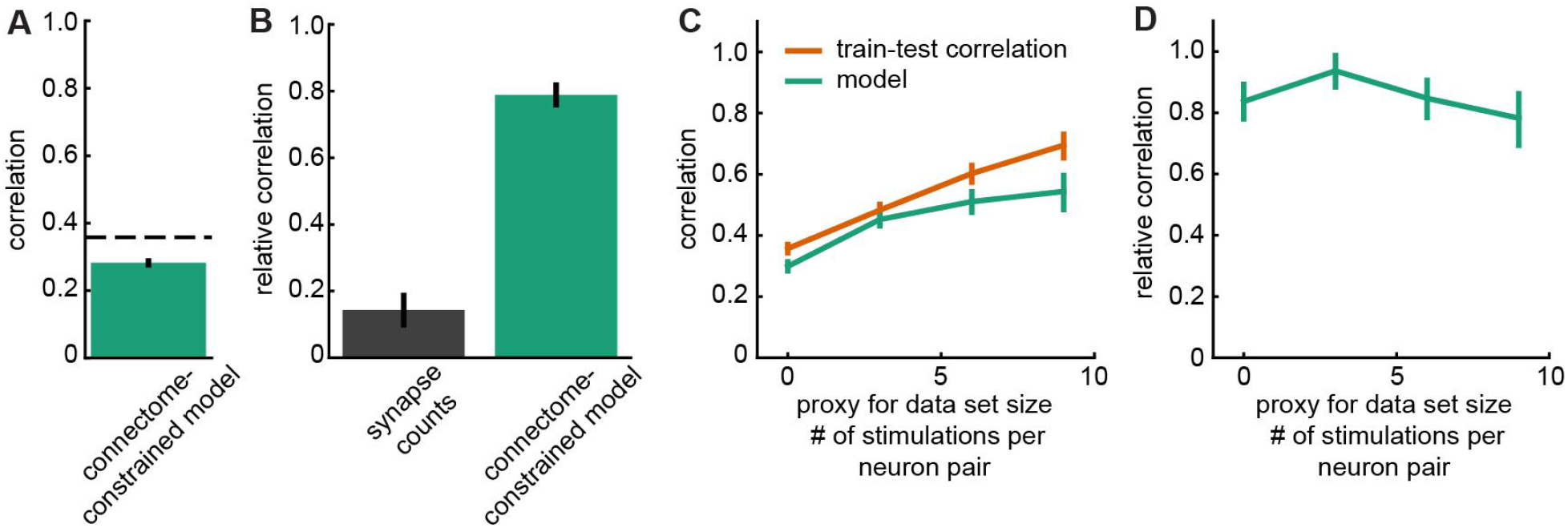
Connectome-constrained model can reproduce the correlative structure of whole-brain data. **A**) Correlation of the model’s matrix of correlation coefficients (MCCs) to the measured MCCs across all neuron pairs (green bar). Dotted line is the train-test correlation for the MCCs. **B**) Relative correlation of the synapse counts and connectome-constrained model to the measured MCCs. Synapse counts are calculated as the sum of the electrical and chemical synapses for each pair of neurons and because they lack sign information they are compared to the absolute value of the MCCs. **C**) Correlation between model MCCs and measured MCCs where the MCCs are segmented into pairs of neurons with different numbers of recording observations as a proxy for data set size. **D**) Relative correlation of the model prediction as a function of data set size. Error bars are 95% confidence intervals, estimated with bootstrapping.

**Supplementary Figure 2.**
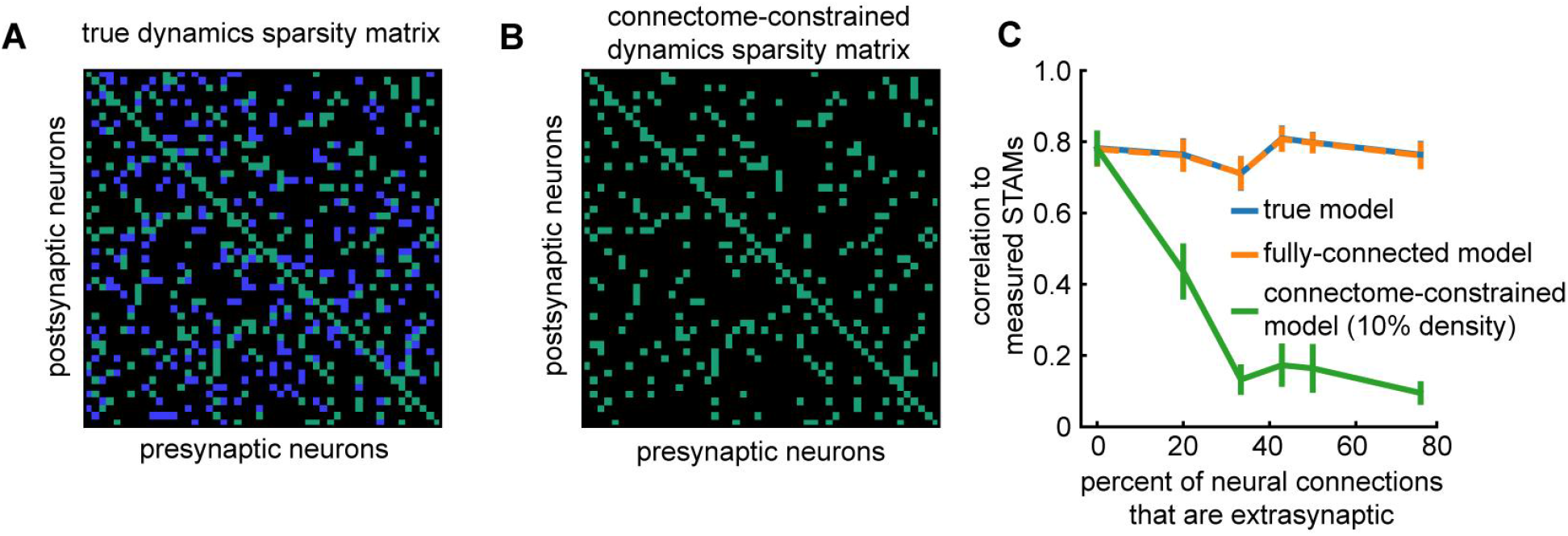
Simulations illustrating that a connectome-constrained model fails to capture synthetic data with significant extrasynaptic signaling. To simulate a system with extrasynaptic signaling, synthetic data was generated from a linear dynamical system with a sparsity constraint on the dynamics weights *W* (example in **A**). Then a second model was trained to reproduce this synthetic data, but using only a subset of the weights of the original model (example in **B**). In this way, the shared weights represent synthetic anatomical weights and the weights that are not shared represent extrasynaptic connections not captured in the synthetic connectome. **A**) Example sparsity constraint of the dynamics weights for the model the data was produced from. Weights with black dots are set to 0. Green and blue dots are randomly drawn from a Gaussian distribution. Green dots are synthetic anatomical connections between a pair of neurons and are shared with the connectome-constrained model in **B**. Blue dots indicate synthetic extrasynaptic connections between a pair of neurons. For this example, 10% of connections are anatomical and 10% are extrasynaptic. **B**) Example sparsity map of the dynamics weights for a connectome-constrained model trained on this synthetic data set. Each connection represents an anatomical connection and is shared with **A. C**) Performance of the models trained on synthetic datasets generated with increasing numbers of extrasynaptic connections. Lines and error bars are mean and standard error of the mean calculated from 10 initializations with different connectivity patterns.

**Supplementary Figure 3.**
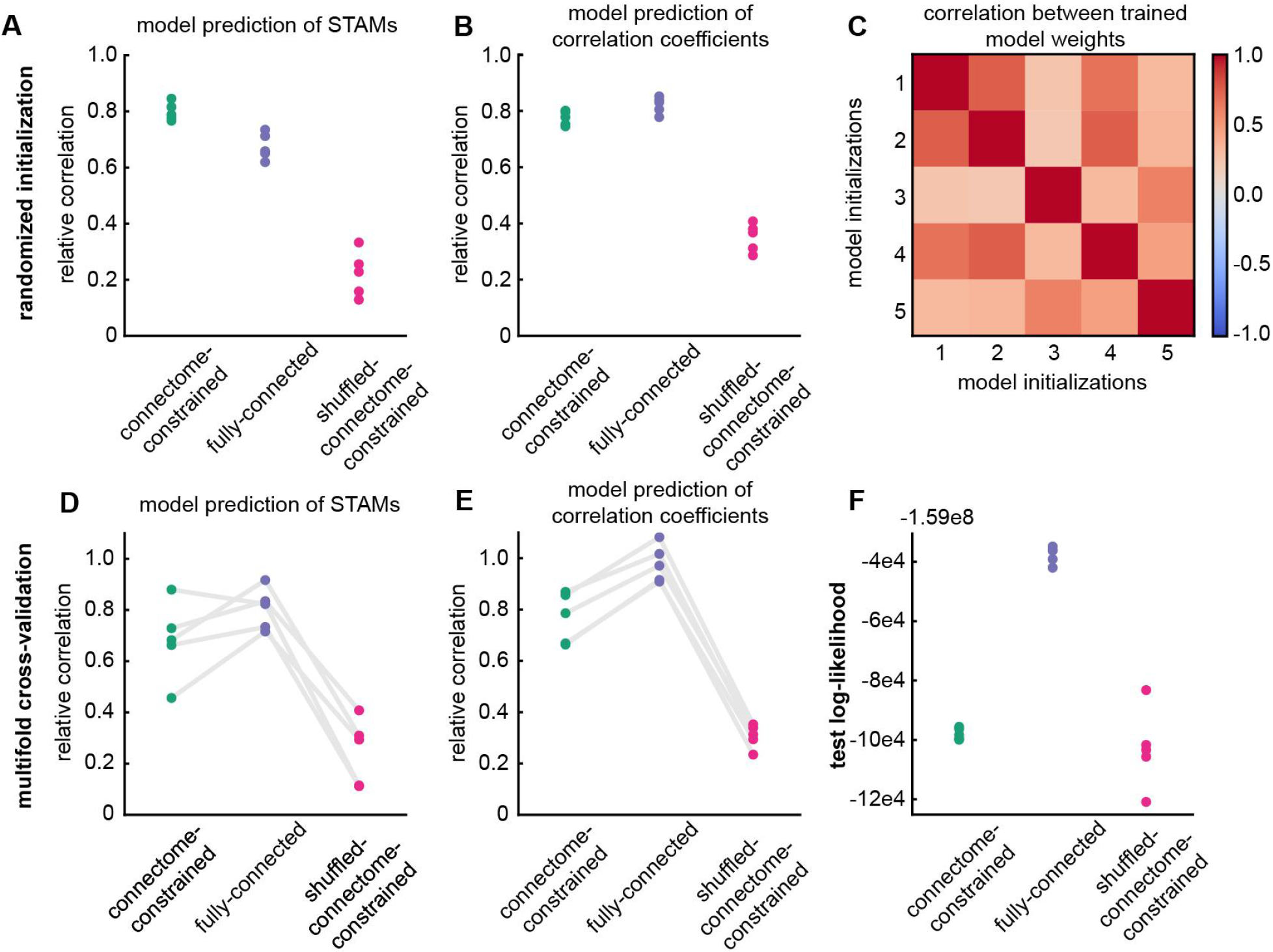
Model performance is insensitive to initialization. Each point represents a fully trained model starting from a randomized initial condition. For the shuffled-connectome-constraint this includes a different randomization of the shuffle. **A**) Relative correlation of model STAMs to measured STAMs across many different initializations. **B**) Relative correlation of model correlation coefficients to measured correlation coefficients across many different initializations. **C**) Correlation between the trained dynamics weights for the connectome-constrained model across the 5 repeated initializations in **AB. DE**) The data set of 110 animals was split into 5 different subsets of 55 animals in the train set and 55 animals in the test set. The model was trained on these 5 different train sets and evaluated on the corresponding test set. **D**) The model’s accuracy in predicting the test set STAMs. **E**) The model’s accuracy in predicting the test set matrix of correlation coefficients. Lines link models trained on the same split of the data. **F**) Test log-likelihood of each model over multiple initializations from **AB**.

**Supplementary Figure 4.**
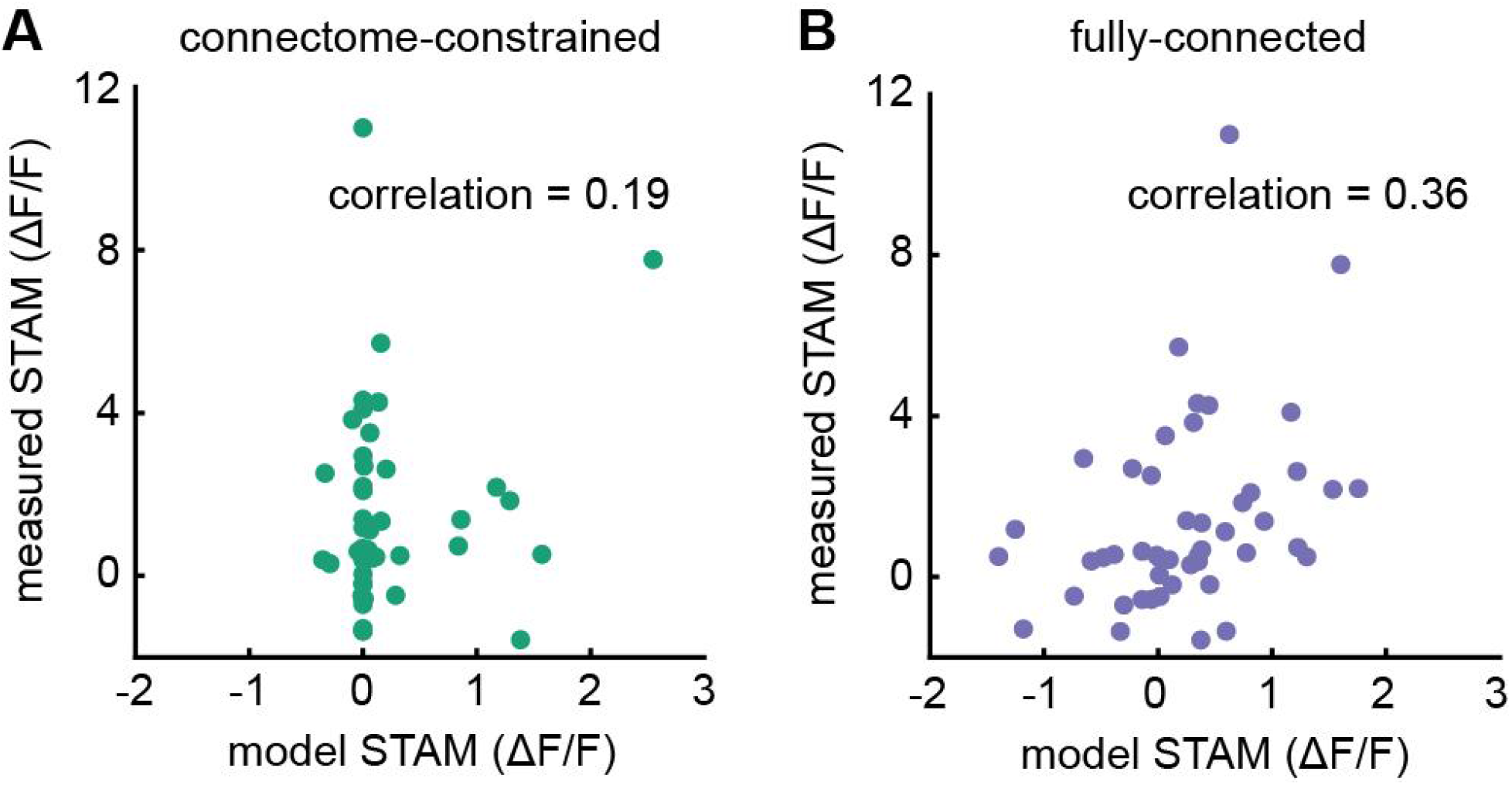
Fully-connected model outperforms connectome-constrained model on putative extrasynaptic connections. **A**) Connectome-constrained model STAM predictions plotted against measured STAMs for 56 putative extrasynaptic connections identified in previous work^2^. **B**) Fully-connected model STAM predictions plotted against the same 56 measured STAMs.

**Supplementary Figure 5.**
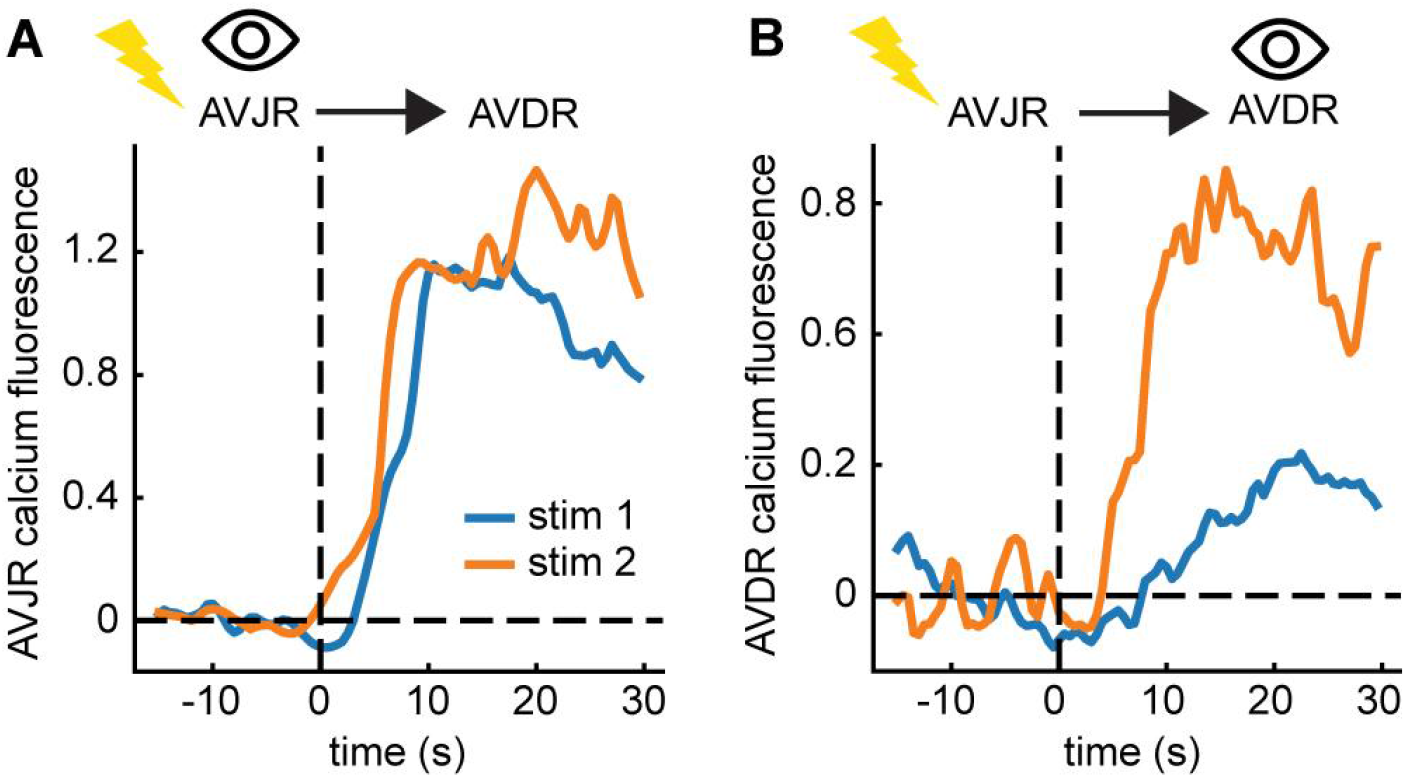
Perturbation responses show variable downstream responses. **A**) Calcium activity in AVJR to two different optogenetic activations of AVJR. Note that AVJR reaches similar levels of calcium activity in both events. **B**) Calcium activity in AVDR during the same 2 stimulation events of AVJR as seen in **A**. Despite the same level of activity in AVJR, AVDR shows variable responses post stimulation.

